# Activation of ACC synthase 2/6 increases stomatal density and cluster on the *Arabidopsis* leaf epidermis during drought

**DOI:** 10.1101/2021.04.27.441570

**Authors:** Ming-zhu Jia, Ling-yun Liu, Chen Geng, Chun-peng Song, Jing Jiang

## Abstract

It is known that the transcription factor SPEECHLESS (SPCH) drives entry of epidermal cells into stomatal lineage, and that the activation of subtilisin-like protease SDD1 reduces stomatal density and cluster on the epidermis. However, there is still a big gap in our understanding of the relationship between stomatal development and the establishment of stomatal density and pattern, especially during drought. Interestingly, 1-aminocyclopropane-1-carboxylic acid (ACC) not only promotes stomatal development, but also is involved in the establishment of stomatal density and pattern. ACC generation comes from the activity of ACC synthase (ACS), while ACS activity could be mediated by drought. This work showed that the Arabidopsis SPCH activated *ACS2/6* expression and ACC-dependent stomatal generation with an increase of stomatal density and cluster under drought conditions; and the possible mechanisms were that ACC-induced Ca^2+^ shortage in stomatal lineage reduced the inhibition of the transcription factor GT-2 Like 1 (GTL1) on *SDD1* expression. These suggest that ACS2/6-dependent ACC accumulation integrated stomatal development with the establishment of stomatal density and pattern by mediating Ca^2+^ levels in stomatal lineage cells on the leaf epidermis, and this integration is directly related to the growth or survival of plants under escalated drought stress.

**Highlight:** ACC synthase ACS2/6 activation integrated stomatal individual development with space setting between stomata by mediating Ca^2+^ levels in stomatal lineage on the leaf epidermis in response to drought.

## Introduction

Stomata are pore structures surrounded by guard cells (GCs) on the leaf epidermis that regulate the exchange of gases (i.e., H_2_O, CO_2_, or O_2_) between plants and the environment (Acharya and Assmann, 2009; Zoulias *et al*., 2018). In evolution from aquatic to terrestrial, plants had generated stomata on epidermis of aerial part to facilitate transpiration (pulling water to shoot), and to guarantee plant survival and life on land (Croxdale, 2000; van Veen and Sasidharan, 2021). For terrestrial plant, stomata can sense environmental water status, especially water deficit or drought, and regulate water loss from plant (Acharya and Assmann, 2009; Zoulias *et al*., 2018). The ability of stomata to regulate water loss is generally estimated from stomatal density (number of stomata per unit leaf area) and pattern (whether stomata are distributed singly or in clusters). Stomatal density and pattern are the consequences of stomatal space setting; for example, stomatal cluster is formed by directly contacting stomata with no intervening epidermal cell, or, alternatively, zero-space establishment (Von Groll *et al*., 2002; Acharya and Assmann, 2009; Zoulias *et al*., 2018). The establishment of stomatal space is, therefore, an important aspect of plant growth and survival under drought conditions (Hepworth *et al*., 2015).

The regulation of stomatal development, which undergirds stomatal spacing, has been extensively investigated. Studies on the model plant *Arabidopsis thaliana* (L.) have shown that stomatal development includes a series of epidermal cell divisions, in which several basic helix-loop-helix transcription factors, such as SPCH, MUTE and FAMA, are involved in this process. SPCH initiates stomatal development to transform meristemoid mother cells (MMCs) into a meristemoids and a sister cell; MUTE converts a sister cell into the guard mother cells (GMCs); then FAMA drives GMCs to form a stoma with differentiated GCs (Hamanishi *et al*., 2012; Zoulias *et al*., 2018). These findings show clearly that SPCH dominates stomatal development. This view is better interpreted, for example, the loss-of-function *spch-1* or *spch-3* homozygous mutant do not produce any stomata (MacAlister *et al*., 2007; Pillitteri *et al*., 2007; Han and Torii, 2016). Furthermore, SPCH expression is known to implicate the establishment of stomatal density (Tripathi *et al*., 2016; Zoulias *et al*., 2018). However, it is still unclear how SPCH integrates stomatal developing with space setting.

Evidences suggest that the subtilisin-like protease Stomatal Density and Distribution 1 (SDD1) participates in space setting between stomata on leaf epidermis, or in the establishment of stomatal density and pattern (Berger and Altmann, 2000; Casson and Gray, 2008; Serna, 2009; Zoulias *et al*., 2018), as evidenced by the following: a loss-of-function *sdd1-1* mutant showed a 2- to 4-fold increase in stomatal density and stomatal clustering in all aerial parts, whereas transgenic *SDD1-*overexpressing plants exhibited a 2- to 3-fold decrease in stomatal density and arrested stomata (Berger and Altmann, 2000; Von Groll *et al*., 2002). In line with these findings, *SDD1*-overexpressing plants displayed diminished transpiration because of a ~25% reduction in abaxial stomatal density or clustering (Yoo *et al*., 2010). Evidently, SDD1 activity is negatively correlated with stomatal density and stomatal clustering ratio. The regulatory mechanism of SDD1 activity has been uncovered. Significantly, the trihelix transcription factor GT-2 LIKE 1 (GTL1) binds to the promoter of the *SDD1* gene and inhibits its expression (Yoo *et al*., 2010; Weng *et al*., 2012; Virdi *et al*., 2015). This inhibition can be relieved by Ca^2+^ increase because Ca^2+^-loaded calmodulin (Ca^2+^-CaM) destabilizes the docking of GTL1 protein to the *SDD1* promoter (Yoo *et al*., 2019). These findings suggest that local Ca^2+^ levels are positively correlated with SDD1 activity as well as stomatal density and the rate of stomatal clustering. Nevertheless, the manner by which SPCH signals information on stomatal development to GTL1-controlled *SDD1* expression or stomatal space setting remains unclear.

The non-proteinogenic amino acid 1-aminocyclopropane-1-carboxylate (ACC) has recently been shown to independently promote stomatal generation by facilitating the differentiation of GMCs into GCs in *Arabidopsis* leaves (Yin *et al*., 2019). Unexpectedly, ethylene is not involved in this process (Yin *et al*., 2019) even though ACC is the precursor of ethylene (Bleecker and Kende, 2000). In fact, ACC is known to be involved in stomatal development and spacing (Acharya and Assmann, 2009). For example, ACC treatments increased the number of stomata by ~33% on the hypocotyl or cotyledon epidermis in *Arabidopsis* (Saibo *et al*., 2003), and also induced stomatal clustering (Serna and Fenoll 1997; Berger and Altmann, 2000). The production of ACC *in vivo* depends on the activity of ACC synthase (ACS), which converts S-adenosylmethionine to ACC (Bleecker and Kende, 2000). Various pieces of experimental evidence strongly suggest that ACS activity is an important mediator of stomata formation. For example, inhibitors of ACS activity, such as aminoethoxyvinylglycine (AVG) and paclobutrazol (PAC), were shown to abolish stomatal appearance (Serna and Fenoll, 1996; Saibo *et al*., 2003; Yin *et al*., 2019). It is known that ACS is encoded by a multi-gene family (Bleecker and Kende, 2000), and that the activity of ACS family members is unique, overlapping, and spatiotemporally specific (Tsuchisaka and Theologis, 2004; Tsuchisaka *et al*., 2009). The *Arabidopsis* genome contains nine *ACS* genes (*ACS1, ACS2, ACS4–9*, and *ACS11*) that encode authentic enzymes (Tsuchisaka *et al*., 2009). Interestingly, the expression of *ACS* genes is induced by drought (Dubois *et al*., 2017; Dubois *et al*., 2018), which suggests that ACS activity may be involved in the stomata-based drought response in *Arabidopsis*. Strikingly, chromatin immune-precipitation assays have indicated that SPCH may regulate the transcription activity of *ACS2* and *ACS6* genes (Lau *et al*., 2014). Nevertheless, further evidence is needed to clarify how SPCH directs ACS2/6 activity during stomatal development.

In this study, we explored the specific involvement of ACS2/6 activity in the drought tolerance of *Arabidopsis* seedlings. Our results revealed that the T-DNA insertion mutants *acs2-1*, *acs6-1*, and *acs2-1acs6-1* are more tolerant to drought than is the wild-type (WT) control. Subsequent research on the underlying mechanism indicated that SPCH activates the expression of *ACS2*, *ACS6*, and *GTL1* by directly binding to their promoters. ACS2/6-dependent ACC accumulation triggers stomatal development and a Ca^2+^ shortage in stomatal lineage cells, and the latter resulted in the repression of GTL1-controlled SDD1 expression. Stomatal density and cluster on the leaf epidermis are thereby increased, leading to increased seedling wilting and even death under intensified drought.

## Materials and methods

### Plant materials and growth conditions

*Arabidopsis thaliana* (Columbia-0 ecotype) was used as WT. The different *ACS2* expression lines, including mutant *acs2-1* (CS16564) with a T-DNA insertion, *ACS2*-complementation (*ACS2*/*acs2-1*), *ACS2*-overexpression (*ACS2*-OE), and *pACS2*::*ACS2-GUS* lines, have been described previously (Han *et al*., 2019). The T-DNA insertion mutant *acs6-1* (CS16569) was obtained from the Arabidopsis Biological Resource Center (USA). The double mutant *acs2-1acs6-1* was created by crossing *acs2-1* with *acs6-1*. Seeds of *spch-3* mutant with a T-DNA insertion were a friendly gift from Professor Sui-wen Hou (MOE Key Laboratory of Cell Activities and Stress Adaptations, Lanzhou, China). These homozygotes with T-DNA insertion were screened according to the method provided by the Salk Institute (http://signal.salk.edu). Seeds of the point mutant *spch-1* was a friendly gift from Professor Xiao-lan Chen (School of Life Sciences, Yunnan University, China), and was identified by PCR amplification and sequencing of the fragment containing the mutation site. All primers used in this study are listed in the Supplementary Table.

Seeds of the transgenic *pSPCH*::*SPCH*-*GFP* line were a friendly gift from Professor Xiao-lan Chen (School of Life Sciences, Yunnan University, China), and GFP expression was detected by hygromycin screening and measurement of fluorescence in leaves. The seeds of Ca^2+^ sensor NES-YC3.6-expressing line were kindly gifted by Professor Jörg Kudla (Molecular Genetics and Cell Biology of Plants, University of Munich, Germany). NES-YC3.6-expressing *acs2-1acs6-1* line was created by crossing *acs2-1acs6-1* with NES-YC3.6-expressing WT plants. Progeny were selected on kanamycin-containing medium and by measuring fluorescence in leaves. All F_3_ progeny meeting the requirements were used in subsequent experiments.

All seeds were collected and stored under the same conditions. Prior to experiments, seeds were surface-sterilized and sown on Murashige-Skoog medium. After 3 days at 4°C in darkness, plates were transferred to a greenhouse (21 ± 2°C, 70% humidity, 100 μmol m^−2^ s^−1^ light intensity, and a 16-h light/8-h dark photoperiod). After germination and growth for 7 days, young seedlings were transplanted into water-saturated soil. Watering was halted according to the requirements of each specific drought treatment described in this paper.

### Creation of transgenic plants

To generate *ACS6*- and *SPCH*-overexpression lines, the full-length coding sequence (CDS) of *ACS6* or *SPCH* was amplified and cloned into the pSUPER 1300 vector. Each construct was then introduced into *Agrobacterium* strain GV3101 and transformed into the target plants by floral infiltration. The same method was used to generate the transformants described below. To generate *ACS6*-complementation (*ACS6*/*acs6-1*) lines, the promoter and CDS of *ACS6* were cloned into a pCAMBIA1300 vector, which was transformed into *acs6-1* plants. To generate *pACS6*::*ACS6*-*GUS* lines, the *ACS6* promoter fragment and full-length CDS were cloned into the promoter-less GUS expression vector pCAMBIA1391, which was then transformed into WT. To generate *pSDD1*::*SDD1*-*GFP* lines, the promoter fragment and CDS of *SDD1* were cloned into a pCAMBIA1300 vector, which was then introduced into *acs2-1acs6-1* and WT. The T_1_ transgenic plants were selected on hygromycin-containing medium, and the T_3_ progeny were used for subsequent experiments.

### Water loss assay

True leaves were collected from 28-day-old plants following previously described methods (Xie *et al*., 2019). The fresh weight of leaves was determined immediately. Leaves of five plants per line were weighed hourly on an electronic balance (Sartorius, Germany) at room temperature (23°C). Water loss was calculated using the following formula: ((W1–W2)/W1) × 100%, where W1 is the initial leaf fresh weight, and W2 is the leaf weight at a given time point.

### Evaluation of stomatal density and rate of stomatal clustering

Stomatal density and clustering ratio were determined according to previously described methods (de Marcos *et al*., 2017; Qi *et al*., 2019). The 6th fully expanded rosette leaves (count up from cotyledons) were used for analyzing the stomatal phenotype of 28-day-old seedlings. Strips were peeled from leaf abaxial epidermis, fixed on a slide, and photographed under a differential contrast interference microscope (LSM710, Zeiss, Germany). Images were acquired under the 20× objective (0.18 mm^2^). Randomly selected images are shown in figures.

For analyses of stomatal density and clustering rate, 25 plants per line per plant were examined. In all counts, a stoma was considered to have a pair of complete guard cells. Stomatal density was calculated as follows: stomatal density = stomatal number/area (mm^2^). The rate of clustered stomata was calculated as follows: number of clusters/(number of stomata + number of clusters).

### RNA extraction and quantitative real-time polymerase chain reaction analyses

Total RNA was extracted using a plant RNA MIDI kit (Life-Feng, Shanghai, China). First-strand complementary DNA (cDNA) was synthesized with a Reverse Transcription system (Toyobo, Osaka, Japan) and was used as the template for quantitative real-time polymerase chain reaction (qRT-PCR) analyses along with 2× SYBR Green I master mix (Vazyme, Nanjing, China). The qRT-PCR analyses were performed on a Roche 480 real-time PCR system (Roche, Mannheim, Germany). The RNA levels were calculated as described by Livak and Schmittgen (2001). The reference gene was *ACTIN8* (AT1G49240).

### GUS staining

Leaves excised from 21-day-old plants or drought-treated plants were incubated overnight in darkness at 37°C in GUS staining solution (0.1 M sodium phosphate buffer, pH 7.0; 0.05 mM K_3_[Fe(CN)_6_]; 0.05 mM K_4_[Fe(CN)_6_]; 1 mg ml^−1^ X-Gluc (Sigma, USA); and 0.1% Triton X-100). After staining, leaves were de-stained with 75% (v/v) ethanol until the chlorophyll was completely removed. The stained leaves were photographed using a Nikon Coolpix or Canon 760D digital camera. Representative photographs are shown in figures.

### Measurement of ACC content

Leaves from the same line of 21-day-old plants were collected and ground into a powder. A 0.1-mg aliquot of powdered sample was transferred into an Eppendorf tube along with 1 ml ultrapure water. To completely extract ACC from leaf tissue, the sample was further fragmented using an ultrasonic crusher (Branson, Danbury, CT, USA). The supernatant was collected, the pH was adjusted to <4, and impurities were removed using 1 ml chloroform. The supernatant was then passed through a column containing C18 adsorbent (Oasis MCX, 30 μm, 3 cc/60 mg, Waters, Milford, MA, USA). The column was eluted with 1 M ammonia in water, with chromatographic methanol as the solvent. The eluent was evaporated to dryness in a Concentrator Plus evaporator (Eppendorf, Hamburg, Germany) under vacuum at 30°C and then re-suspended in solution (chromatographic methanol: 0.1% (v/v) acetic acid, 1:9). Samples were analyzed using an Applied Biosystems MDS SCIEX 4000 QTRAP liquid chromatography-tandem mass spectrometry system (AB Sciex, Foster City, CA, USA). Standard ACC (Sigma-Aldrich, Steinheim, Germany) was used for the quantitative analysis.

### Protein extraction and western blotting

Leaves were collected according to the experimental requirements and ground into a powder. Powdered samples were transferred to RIPA lysis buffer (Boster Biotechnology, Wuhan, China) and micro-centrifuged at 16,000 *g* for 15 min at 4°C. The concentration of crude protein in the supernatant was determined using a NanoDrop 2000 (Thermo Scientific, Wilmington, DE, USA). The crude protein was separated by 12% SDS-PAGE and then transferred to a nitrocellulose filter membrane (Millipore, Billerica, MA, USA) using a Trans-Blot Semi-Dry transfer cell (Bio-Rad, Hercules, CA, USA). The membrane was then incubated at room temperature for 1-2 h in blocking solution before incubation with anti-GFP mouse monoclonal antibodies (1:10,000; Proteintech, Chicago, IL, USA) for 2 h at room temperature. The membrane was subjected to three 10-min washes with TBST and then incubated overnight at 4°C with horseradish peroxidase-conjugated secondary antibody (Proteintech). Protein bands were detected using a BeyoECL Plus kit (Beyotime, Shanghai, China) and then visualized using a Fusion FX7 Spectra system (Vilber Lourmat, Marne-la-Vallée, France). An anti-GAPDH antibody (1:5000; Proteintech) was used as the loading control.

### Chromatin immunoprecipitation analyses

Immature leaves collected from *pSPCH::SPCH*-*GFP* of 21-day-old plants were cross-linked using 1% formaldehyde under vacuum for 10 min according to the EZ-ChIP chromatin IP kit protocol (Thermo Scientific). After washing with phosphate-buffered saline solution, leaves were ground in liquid nitrogen and then suspended in SDS lysis buffer containing protease inhibitor cocktail. The DNA was sheared into small fragments (300–500 bp). The sheared chromatin was immune-precipitated with GFP antibodies (Proteintech) overnight at 4°C. The ChIP DNA products were analyzed by RT-qPCR using three pairs of primers synthesized to amplify approximately 200-bp DNA fragments of the promoter region of *ACS2* or *ACS6,* which were used in the ChIP analysis. Primers annealing to promoter regions of two *Arabidopsis* genes lacking an SPCH binding site were used as negative controls. An unrelated DNA sequence from the *ACTIN8* gene was used as an internal control.

### Transient transcription dual-luciferase assays

Detection was performed according to previously described methods (Bao *et al*., 2014). The 2400-bp promoter sequence of *ACS2* was divided into 3 fragments (−1 to −1000, −900 to −1600, and −1500 to −2400 bp). The 2600-bp promoter sequence of *ACS6* was also divided into 3 fragments (−1 to −1000, −900 to−2000, and −1900 to −2600 bp), and the −1 to −780 bp promoter sequence of GTL1 was selected. Each fragment was cloned into pGreen II 0800-Luc to construct the corresponding reporter plasmid. The coding sequence of *Arabidopsis SPCH* was cloned into pGreenII 62-SK to construct the 35S-SPCH effector plasmid. The *Agrobacterium* strain GV3101 (pSoup-p19) was incubated in yeast mannitol medium and finally re-suspended in buffer to a final concentration of OD_600_ = 1.0. Equal amounts of different combined bacterial suspensions were infiltrated into young leaves of tobacco plants using a needleless syringe. After 3 days, the infected leaves were sprayed with D-luciferin (sodium salt) (Yeasen, Shanghai, China) and placed in darkness for 5 min. Firefly luciferase (LUC) signals were then detected using the NightSHADE system (LB 985, Berthold Technologies, Bad Wildbad, Germany). The ratio of LUC activity to Renilla luciferase (REN) activity was measured using a Dual-Luciferase Reporter Gene Assay kit (Solarbio, Beijing, China). Briefly, the tobacco leaves were ground in liquid nitrogen, and the extract was incubated in a low-temperature buffer. The LUC/REN ratio was measured using an enzyme standard instrument (Tecan, Männedorf, Switzerland).

### Monitoring of Ca^2+^ levels in stomatal lineage cells

The Ca^2+^ levels in stomatal lineage cells were monitored according to Krebs *et al*. (2012). Immature leaves of *Arabidopsis* seedlings expressing the fluorescence resonance energy transfer (FRET)-based Ca^2+^ sensor NES-YC3.6 (Nagai *et al*., 2004; Krebs *et al*., 2012) were collected from the same position. During confocal laser scanning, strips were peeled from leaf abaxial epidermis and then fixed on a slide on the loading platform. The relative fluorescence intensity of YC3.6 protein was recorded under a Nikon A1 Plus laser scanning confocal microscope (Nikon, Tokyo, Japan) with the following scanning parameters: image dimension = 1024 × 1024, pinhole radius = 38.31 μm, scanning speed = 0.25, zoom = 3×, objective = 60× (water), numerical aperture = 1.27), plan apochromat objective, power = 6% (445 nm solid laser). Images were acquired every 5s. Emissions from cyan fluorescent protein (CFP; 465-499 nm) and FRET-dependent cpVenus (525-555 nm) in stomatal lineage were detected simultaneously. The cpVenus/CFP emission ratio was analyzed using NIS-Elements AR software.

### Statistical analysis

All experiments were independently repeated using three biological replicates and three technical replicates at least. Differences among treatments were compared using Student’s *t*-test. *P*-values < 0.05 (*) and < 0.01 (**) were considered to correspond to significant and extremely significant differences, respectively.

## Results

### ACS2/6 activation and ACC accumulation facilitated water evaporation from leaves in response to drought

Studies have shown that the activity of ACS2 in rice (Zhang *et al*., 2013) and ACS6 in maize (Young *et al*., 2004) regulates seedling sensitivity to drought, and that drought induces ACS2/6 activation and ACC accumulation in *Arabidopsis* (Catalá *et al*., 2014; Dubois *et al*., 2018). Therefore, we examined the possible roles of ACS2 and ACS6 in the drought response of *Arabidopsis* seedlings.

The expressions of *ACS2* and *ACS6* were modified in several genetic materials. For example, compared with WT, the loss-of-function mutant lines *acs2-1*, *acs6-1*, and *acs2-1acs6-1* showed significantly reduced *ACS2* or *ACS6* mRNA levels; the transgenic *ACS2*-OE(#1) and *ACS6*-OE(#1) over-expression lines had significantly elevated *ACS2* and *ACS6* mRNA levels, respectively; the transgenic complemented lines *ACS2*/*acs2-1*(#1) and *ACS6*/*acs6-1*(#1) exhibited no changes in *ACS2* or *ACS6* expression, respectively (Fig. 1A). Interestingly, the expressions of *ACS6* and *ACS2* were relatively unchanged in the single mutants *acs2-1* and *acs6-1*, respectively (Fig. 1B). We first checked the growth phenotypes of the various ACS2/6 expression lines in response to drought. Under normal watering (control) conditions, no significant water-losing phenotypes were apparent among these diverse expression lines. Under gradually intensifying drought caused owing to stopping watering, however, the phenotypes were obviously different. In particular, WT, *ACS2*/*acs2-1*(#1), *ACS6*/*acs6-1*(#1), *ACS2*-OE(#1), and *ACS6*-OE(#1) seedlings withered and some even died after water was withheld for 12 days and the soil water content dropped to ~39%, whereas wilting and drying symptoms were clearly alleviated in the mutants *acs2-1*, *acs6-1*, and *acs2-1acs6-1* seedlings (Supplementary Fig. S1). Preliminarily data showed that ACS2/6 activation promoted dehydration and wilting of seedlings under drought conditions.

**Fig. 1.**
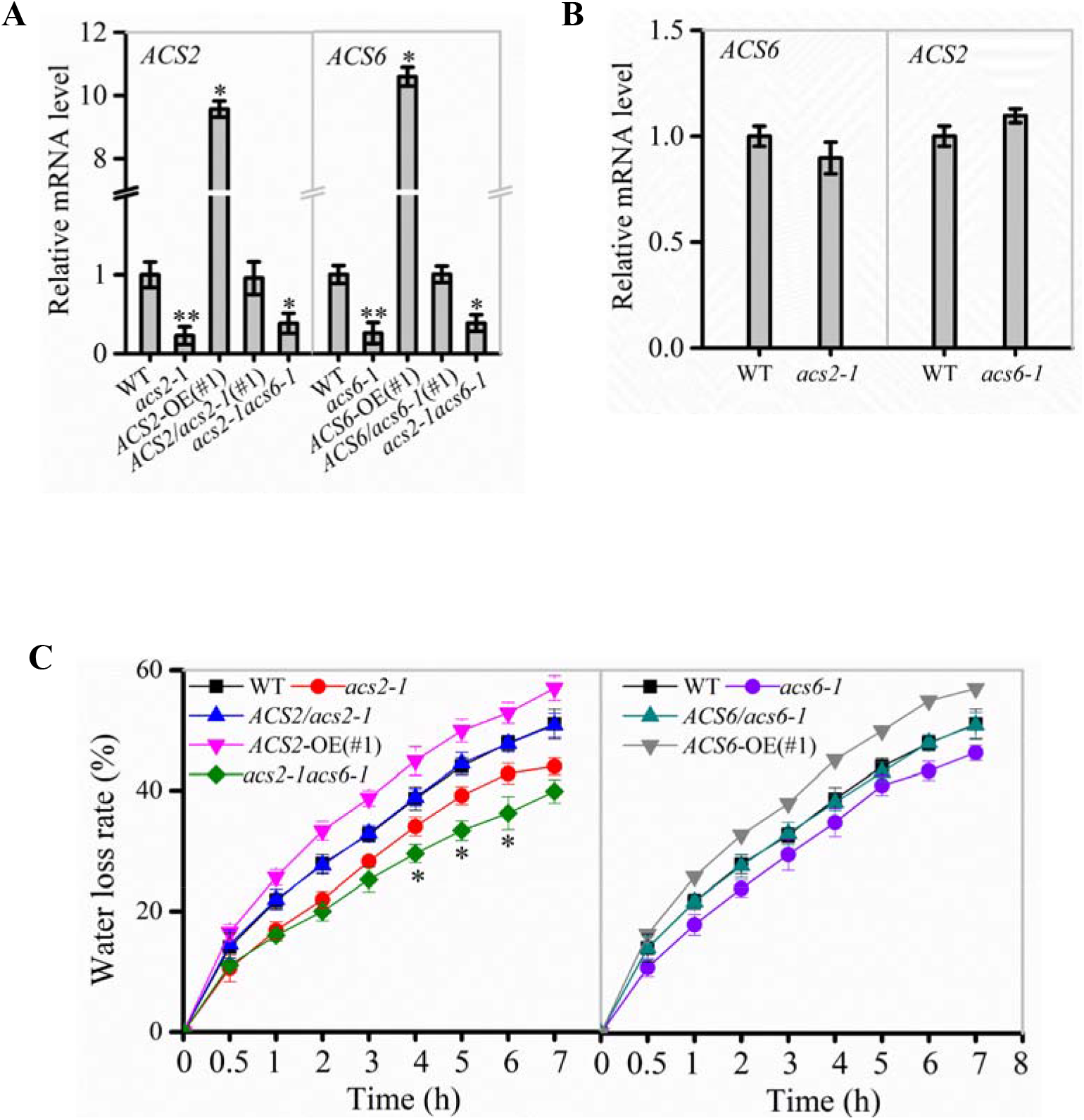
Effect of ACS2/6 expression on water loss from leaves of *Arabidopsis* seedlings. (A) qRT-PCR analysis of *ACS2* and *ACS6* mRNA levels in various lines, including the WT; loss-of-function mutants *acs2-1*, *acs6-1*, and *acs2-1acs6-1*; overexpression lines *ACS2*-OE(#1) and *ACS6*-OE(#1); and complementation lines *ACS2*/*acs2-1*(#1) and *ACS6*/*acs6-1*(#1)*. ACTIN8* was used as a reference gene. Experiments were repeated three times. Values are means ± SD (Student’s *t*-test; *, *P* < 0.05; **, *P* < 0.01). (B) qRT-PCR analysis of *ACS6* mRNA levels in *acs2-1* and *ACS2* mRNA levels in *acs6-1*. Experiments were repeated three times with similar results. (C) Relative rate of water loss over time from detached rosette leaves of 28-day-old plants. All true leaves of five plants of the same line grown under identical conditions were collectively weighed every hour. The data represent the water loss percentage at a given time point, calculated as follows: ((initial weight – weight at each time point) / initial weight) × 100. Experiments were repeated at least three times with similar results. Values are means ± SD (Student’s *t*-test; *, *P* < 0.05).

The rate of water evaporation from leaves of these lines was monitored under drought conditions. The water evaporation rate was decreased in *acs2-1*, *acs6-1*, and *acs2-1acs6-1* leaves, compared with that in WT. This decrease was less pronounced in the double mutant *acs2-1acs6-1* than in the single mutants *acs2-1* and *acs6-1* (Fig. 1C). Conversely, the water evaporation rate was significantly increased in *ACS2*-OE(#1) and *ACS6*-OE(#1) compared with that of WT, whereas no significant change was observed in *ACS2/acs2-1*(#1) or *ACS6*/*acs6-1*(#1) (Fig. 1C). Data suggest that ACS2/6 activation was positively correlated with the rate of water evaporation from leaves under drought conditions.

The characteristics of ACS2/6 expression in leaves were examined in response to drought treatment. Histochemical staining revealed that the higher GUS-marked ACS2/6 expression was in immature leaves, followed by senescent leaves, and then mature leaves in WT plants; After withholding water for 6 days, ACS2 and ACS6 expressions in WT were significantly increased in immature leaves, slightly increased in mature leaves, and unchanged in senescent leaves (Fig. 2A). Meantime, quantitative PCR showed the same results (Fig. 2B). This is, ACS2 and ACS6 expressions were always higher in senescent leaves regardless of drought, whereas both expressions increased in response to drought in non-senescent leaves. Next, ACS2/6-dependent ACC accumulation was analyzed in leaves. Under normal conditions, ACC mainly accumulated in immature and senescent leaves of WT seedlings, whereas ACC accumulated primarily in immature leaves in response to a 6-day halt in watering. More specifically, ACC levels were, respectively, 1.94- and 1.33-times higher in immature and mature leaves of WT after withholding water (Fig. 2C). In contrast to WT, the double mutant *acs2-1acs6-1* did not significantly accumulate ACC in the leaves in response to drought (Fig. 2C). These data indicate that drought induced ACS2/6 activation and ACC accumulation in non-senescent leaves of *Arabidopsis* seedlings.

**Fig. 2.**
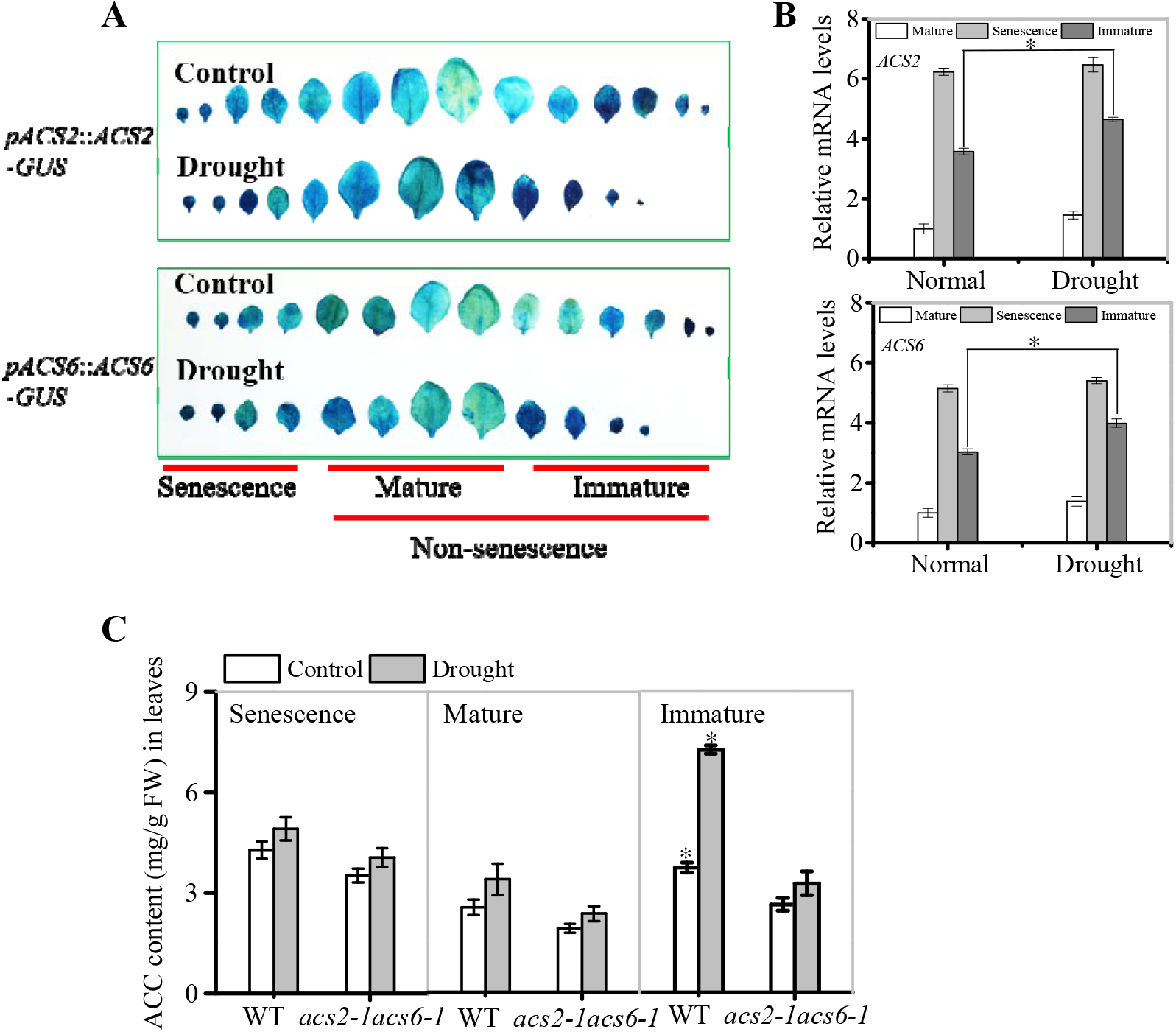
Effects of drought on ACS2 and ACS6 expressions and ACC accumulation in leaves of different ages. (A) ACS2 or ACS6 expression was monitored in immature, mature and senescence leaves of recombinant *pACS2*::*ACS2-GUS* or *pACS6*::*ACS6-GUS* lines, respectively, with or without drought treatment. Representative images are shown. Experiments were repeated three times with similar results. (B) ACS2 or ACS6 expression was monitored by qPCR in immature, mature and senescence leaves of WT, respectively, with or without drought treatment. Representative images are shown. Experiments were repeated three times with similar results. Values are means ± SD (Student’s *t*-test; *, *P* < 0.05). (C) HPLC analysis of ACC accumulation in leaves of 21-day-old seedlings with or without drought treatment. Experiments were repeated three times with similar results. Values are means ± SD (Student’s *t*-test; *, *P* < 0.05).

### ACS2/6 activation affected stomatal density and pattern on leaf abaxial epidermis

Evidences suggest that the ACS activity was required for stomatal development (Serna and Fenoll, 1996; Saibo *et al*., 2003; Young *et al*., 2004; Zhang *et al*., 2013; Lau *et al*., 2014; Yin *et al*., 2019), we thus analyzed the effects of ACS2/6 activity on stomatal density and pattern on the leaf abaxial epidermis.

Images of stomata on the abaxial epidermis of the 6th (count up from cotyledon) mature leaves of the various ACS2/6 expression lines under normal and drought conditions are shown in Fig. 3A. Under normal watering conditions, stomatal density and the percentage of clustering were slightly decreased in the mutants *acs2-1*, *acs6-1*, and *acs2-1acs6-1*, but slightly increased in *ACS2*-OE(#1) and *ACS6*-OE(#1), compared with that in WT (Fig. 3B). After halting watering for 6 days, however, stomatal density was significantly reduced in *acs2-1* (177.8 ± 8.2 mm^−2^), *acs6-1* (183.3 ± 6.2 mm^−2^), and *acs2-1acs6-1* (161.1 ± 8.2 mm^−2^), but significantly increased in *ACS2*-OE(#1) (255.6 ± 6.9 mm^−2^) and *ACS6*-OE(#1) (261.1 ± 4.9 mm^−2^), compared with that in WT (205.6 ± 5.4 mm^−2^) (Fig. 3B). The percentage of the pairs of directly contacting stomata was significantly higher in *ACS2*-OE(#1) (~ 3.1%) and *ACS6*-OE(#1) (~ 2.7%) than in WT (~ 0.2%). In contrast, the stomatal clustering rates were lower in the mutants *acs2-1* (~ 0.12%), *acs6-1* (~ 0.15%) and *acs2-1acs6-1* (~ 0.1%) than in WT (Fig. 3C). Evidently, ACS2/6 activation increased stomatal density and cluster on the abaxial epidermis. In addition, application of 5–10 μM ACC also significantly increased the stomatal density and clustered stomata on the leaf abaxial epidermis of WT seedlings (Supplementary Fig. S2). This validation experiments indicate that appropriate concentrations of ACC increased stomatal density and cluster on leaf epidermis.

**Fig. 3.**
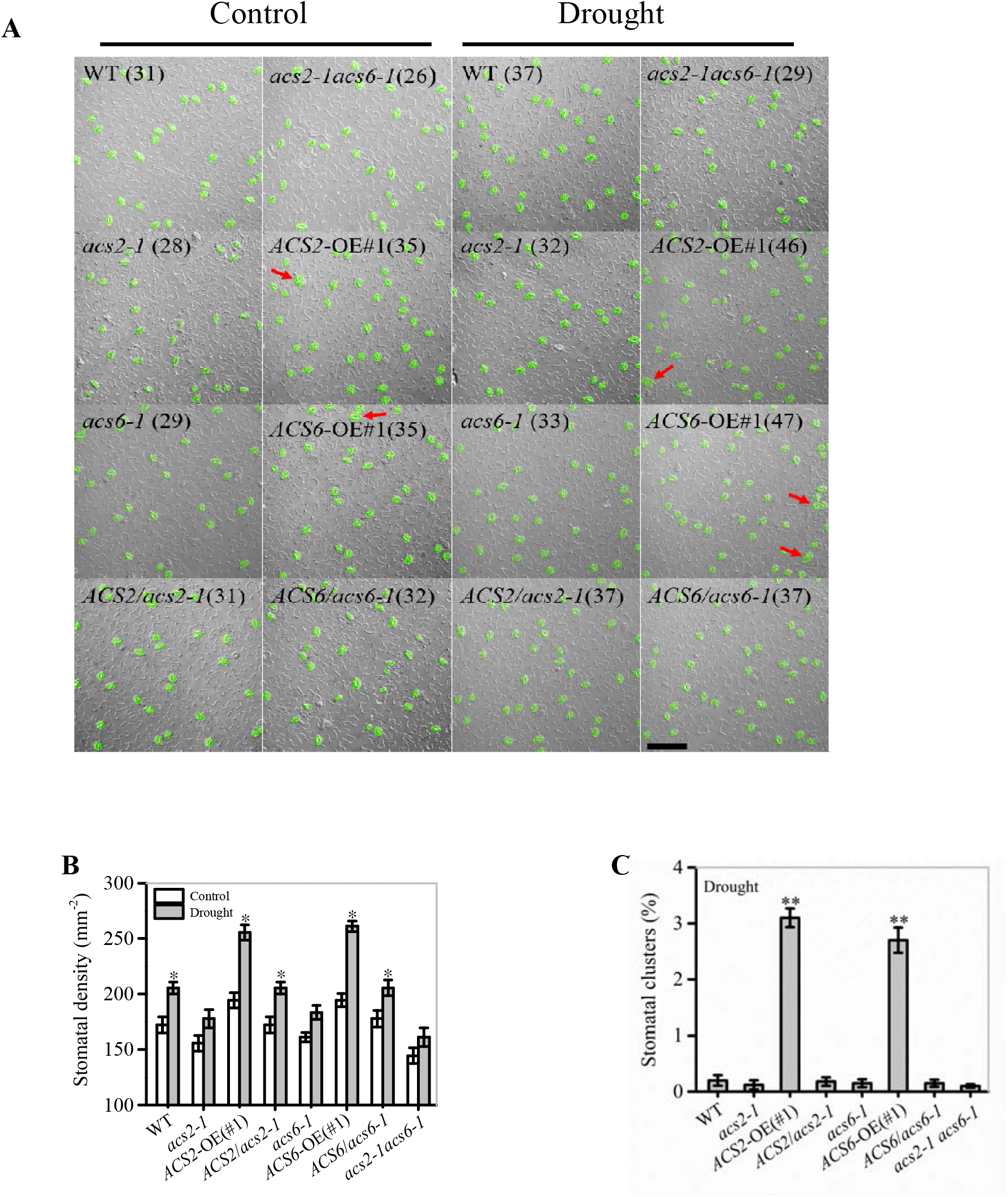
Correlation between ACS2/6 activation and stomatal density and rate of stomatal clustering. (A) Images of stomata distributed on strips of leaf abaxial epidermis under differential interference contrast (DIC) microscopy. Stomata and stomatal clusters are colored in green, and stomatal clusters are indicated by red arrows. The number of stomata is given in parentheses in each image. 6th rosette were collected under drought or normal watering conditions from 28-day-old seedlings of the WT control; mutants *acs2-1*, *acs6-1*, and *acs2-1acs6-1*, overexpression lines *ACS2*-OE(#1) and *ACS6*-OE(#1), and complementation lines *ACS2*/*acs2-1*(#1) and *ACS6*/*acs6-1*(#1). Experiments were performed three times with similar results. The black scale bar represents 100 μm. (B) and (C) Statistical analysis of stomatal density (B) and percentage of stomatal clusters (C). Stomata and stomatal clusters on 25 leaves of 25 seedlings were counted. Values are means ± SD (Student’s *t*-test; *, *P* < 0.05; **, *P* < 0.01).

### SPCH could bind to promoters of *ACS2/6* and regulate their expression

The above data suggest that the effect of ACS2/6-dependnet ACC accumulation is similar to that of SPCH (Tripathi *et al*., 2016; Zoulias *et al*., 2018) in promoting stomatal density on the leaf epidermis, we speculated that SPCH may mediate ACS2/6 expression activity. Although a profile list generated by genome-wide ChIP-based sequencing of the targets of SPCH included both ACS2 and ACS6 (Lau *et al*., 2014), direct experimental evidence was still lacking.

To explore whether SPCH affect *ACS2* and *ACS6* expression, we checked mRNA levels of *ACS2* and *ACS6* in the loss-of-function *spch-1* and *spch-3* mutant seedlings, respectively. In order to explore the stomatal development on the epidermis of true leaves, the heterozygote of *spch-1* and *spch-3* were used, because the two homozygotes cannot grow true leaf (MacAlister *et al*., 2007; Pillitteri *et al*., 2007; Han and Torii, 2016). Observations indicated that both *spch-1* and *spch-3* had significantly reduced *ACS2* and *ACS6* mRNA levels, respectively, whereas *SPCH*-OE lines had significantly increased mRNA levels, compared with the WT control (Fig. 4A). Interestingly, drought similarly induced the expressions of *SPCH*, *ACS2*, and *ACS6* genes (Fig. 4B). Data implies that SPCH activity was positively correlated with the expression of *ACS2* and *ACS6*.

**Fig. 4.**
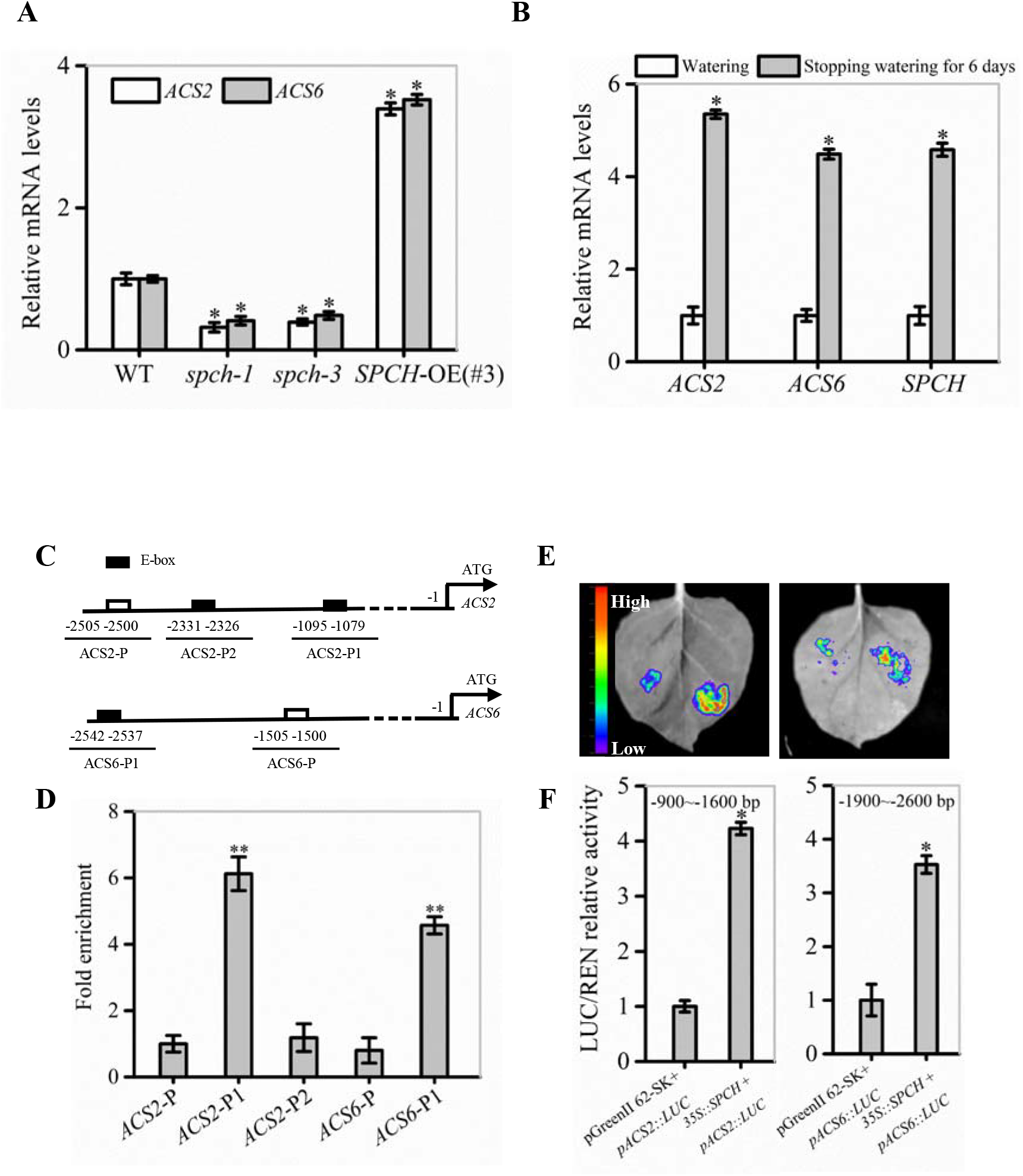
Evidence for the function of SPCH as a transcription factor of *ACS2* and *ACS6* genes. (A) Levels of *ACS2* and *ACS6* mRNA in immature leaves of WT control, loss-of-function *spch-1* and *spch-3* mutant, and *SPCH*-overexpressing *SPCH*-OE(#3) plants based on qRT-PCR. Experiments were repeated three times with similar results. Values are means ± SD (Student’s *t*-test; *, *P* < 0.05). (B) Relative levels of *ACS2*, *ACS6*, and *SPCH* mRNA transcripts in leaves of 21-day-old WT seedlings under normal watering conditions or after 6 days without watering. Experiments were repeated three times with consistent results. Values are means ± SD (Student’s *t*-test; *, *P* < 0.05). (C) Diagram of the relative position of E-boxes (CACGTG or CGCGTG) and a reference DNA region in *ACS2* and *ACS6* gene promoters. Black rectangles indicate E-boxes in *ACS2* (ACS2-P1 and ACS2-P2) and *ACS6* (ACS6-P1) promoter regions, while white rectangles represent reference regions, namely randomly selected DNA fragments from *ACS2* (ACS2-P) and *ACS6* (ACS6-P) promoter regions. (D) Relative abundance of SPCH-immunoprecipitated DNA fragments as determined by qRT-PCR. All experiments, which included three biological replicates, gave similar results. Values are means ± SD (Student’s *t*-test; **, *P* < 0.01). (E) and (F) Binding of SPCH protein to *ACS2* and *ACS6* genes in tobacco leaves in a transient transcription dual-luciferase assay. The size and intensity of LUC fluorescence signals recorded by IndiGO software are proportional to binding ability (E). Relative binding ability was evaluated quantitatively by calculating the ratio of the fluorescence intensity of firefly luciferase (LUC) to that of an internal control, Renilla luciferase (REN) (F). Values are means ± SD (*n* = 3). Asterisks indicate significant differences (*, *P* < 0.05) compared with leaf regions injected with *Agrobacterium* harboring an empty vector.

To confirm this experimentally, we used ChIP assays to detect the interaction between the transcription factor SPCH and the promoters of *ACS2* and *ACS6*. The *in silico* analyses revealed three E-box motifs in the 3.0-kb promoter region of the *ACS2* gene: CGCGTG and CACGTG (at −1079 and −1090), collectively named ACS2-P1 because of their close proximity, and CACGTG (at −2326), designated as ACS2-P2. Only one E-box motif was present in the 3.0-kb promoter region of the *ACS6* gene: CACGTG (at −2537), named ACS6-P1 (Fig. 4C). After randomly selecting DNA fragments from their promoter regions with the same length as the E-boxes (named ACS2-P and ACS6-P) as the reference, ChIP assays were performed to measure levels of immune-precipitated DNA fragments by SPCH protein *in vivo*. In these assays, the abundance of DNA fragments from *ACS2* promoters ACS2-P1 and ACS2-P2 was, respectively, 6.13- and 1.18-fold higher than that of the control ACS2-P (Fig. 4D). Similarly, the abundance of ACS6-P1 immune-precipitated by SPCH protein was 4.57-fold higher than that of the control ACS6-P (Fig. 4D). Next, we conducted transient transcription activity assays to verify the binding of SPCH to the promoters of *ACS2* and *ACS6*. According to the results, the fluorescence intensity of LUC linked to the specific promoter fragment of *ACS2* (−900 to −1600 bp, containing ACS2-P1) was increased in the presence of SPCH, with LUC activity 4.2-times higher than that of the blank LUC control (Fig. 4E, F). Similarly, the activity of LUC linked to the promoter fragment of *ACS6* (−1900 to −2600 bp, containing ACS6-P1) in the presence of SPCH was 3.7-times higher than that of the blank LUC control (Fig. 4E, F). This stimulatory effect was specific, as SPCH did not induce LUC activity alone or when linked to E-box-free promoter fragments of *ACS2* or *ACS6* (Supplementary Fig. S3). In other words, SPCH directed the transcription of *ACS2* or *ACS6* by docking to each of their promoter regions.

To verify that SPCH promoted *ACS2/6* expression, we monitored the effects of SPCH activity on ACC levels. The results showed that the SPCH-overexpressing lines *SPCH*-OE(#1), *SPCH*-OE(#2), and *SPCH*-OE(#3) had significantly increased ACC levels (Fig. 5A), but the mutants *spch-1* or *spch-3* had reduced ACC levels in immature leaves, as compared with the ACC levels in leaves of WT (Fig. 5B). Evidently, SPCH-directed *ACS2/6* expression was directly related to ACC accumulation in immature leaves. Further observations helped to explain how SPCH mediated stomatal development *via ACS2/6-*dependent ACC production. The single mutant *spch-1*, the double mutant *acs2-1acs6-1*, and the triple mutant *spch-1acs2-1acs6-1* had significantly reduced stomatal densities under normal or drought conditions, compared with WT (Fig. 5C, D). In addition, the stomatal density on leaves was lower in the triple mutant *spch-1acs2-1acs6-1* than in its parents *spch-1* and *acs2-1acs6-1* (Fig. 5C, D). Notably, ACS2- or ACS6-overexpression in *spch-1* reversed the reduction in stomatal density (Fig. 5D). These observations suggest that SPCH activity induced ACS2/6 activation and ACC accumulation.

**Fig. 5.**
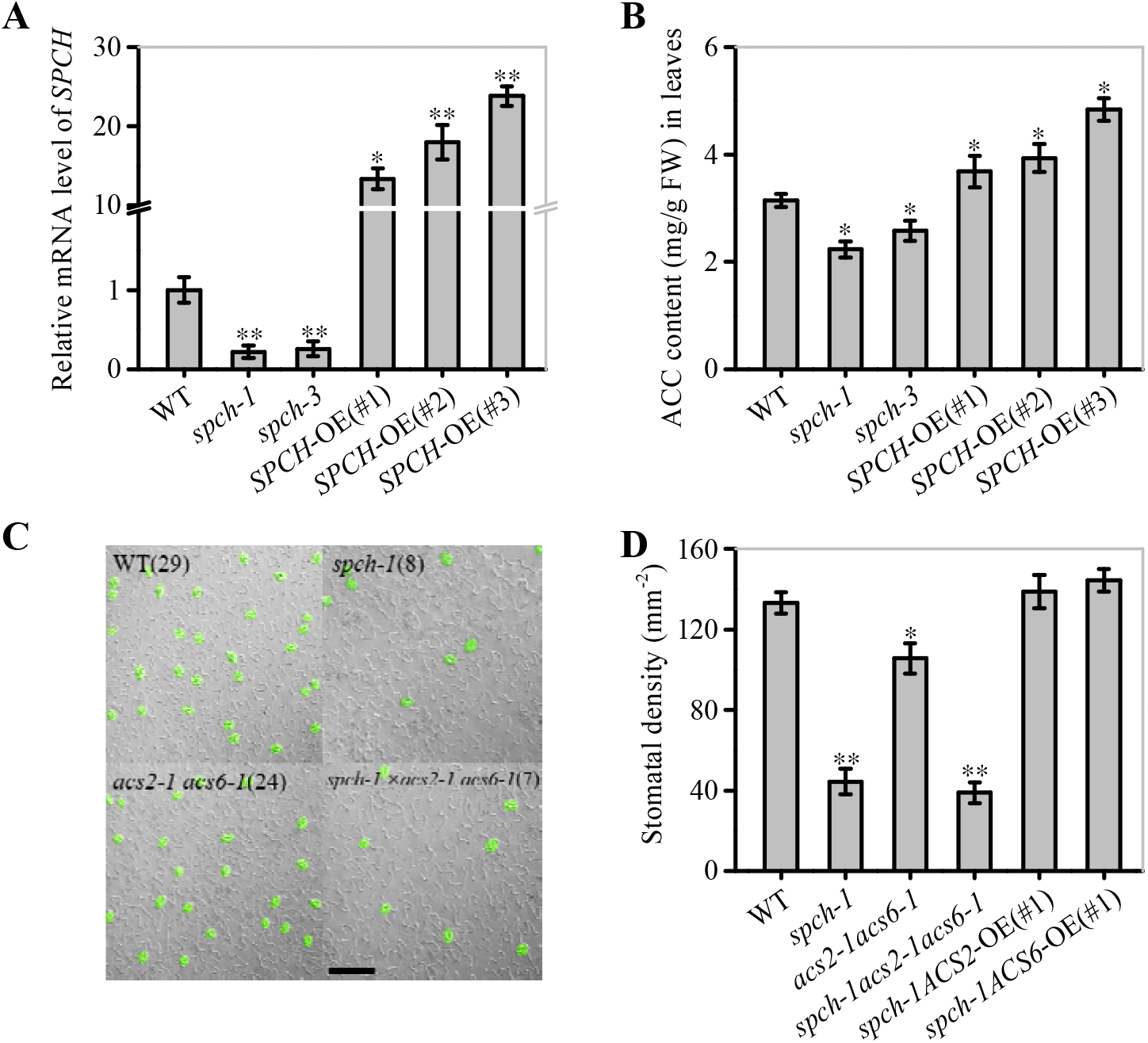
Correlation between SPCH activity and ACS2/6-dependent ACC accumulation and stomatal density and pattern. (A) and (B) qRT-PCR-based relative levels of *SPCH* mRNA transcripts and ACC content in immature leaves of WT control, loss-of-function mutant *spch-1* and *spch-3*, and *SPCH*-overexpressing *SPCH*-OE(#1), *SPCH*-OE(#2), and *SPCH*-OE(#3) plants. The *ACTIN8* gene was used as an internal control. The experiment was repeated three times with consistent results. Values are means ± SD (Student’s *t*-test; *, *P* < 0.05; **, *P* < 0.01). (C) and (D) DIC images and statistical summary of stomatal density and patterning on the abaxial epidermis of *spch-1*, *acs2-1acs6-1*, and *spch-1acs2-1acs6-1* plants. Numbers of stomata are indicated in parentheses in each image. Stomata are false colored in green. The black scale bar represents 100 μm (C). Stomata on 25 leaves of 25 seedlings were counted (D). Values are means ± SD. Significant differences are indicated by asterisks (Student’s *t*-test; *, *P* < 0.05; **, *P* < 0.01).

### ACC accumulation decreased SDD1 expression levels in leaves

Evidences have shown that SDD1 expression reduces stomatal density and cluster (Yoo *et al*., 2010; Yoo *et al*., 2019), but exogenously applied ACC increases stomatal density and cluster (Serna and Fenoll, 1996; Saibo *et al*., 2003; Acharya and Assmann, 2009). We therefore verified whether ACC cooperates with SDD1 to establish stomatal density and cluster.

The effect of ACS2/6-dependent ACC accumulation on SDD1 expression was surveyed in immature leaves. The *SDD1* mRNA levels in *acs2-1*, *acs6-1*, *acs2-1acs6-1*, *ACS2*-OE(#1), and *ACS6*-OE(#1) were, respectively, 1.88-, 1.89-, 2.18-, 0.64-, and 0.67-fold that in WT (Fig. 6A). This result suggests that *SDD1* expression was negatively correlated with ACS2/6 activation in immature leaves. Further monitoring of SDD1 protein levels by western blotting indicated that GFP-marked SDD1 protein levels in immature leaves were higher in *acs2-1acs6-1* than in WT under drought conditions (Fig. 6B). Likewise, ACC treatment reduced protein levels of SDD1 in WT leaves compared with the blank control (Fig. 6B). This result is consistent with the expectation that ACC treatment would reduce *SDD1* mRNA transcript levels in immature leaves of WT (Supplementary Fig. S4). These observations suggest that ACS2/6-generated ACC impeded *SDD1* gene expression and SDD1 protein levels, thereby increasing stomatal density and cluster on the leaf epidermis.

**Fig. 6.**
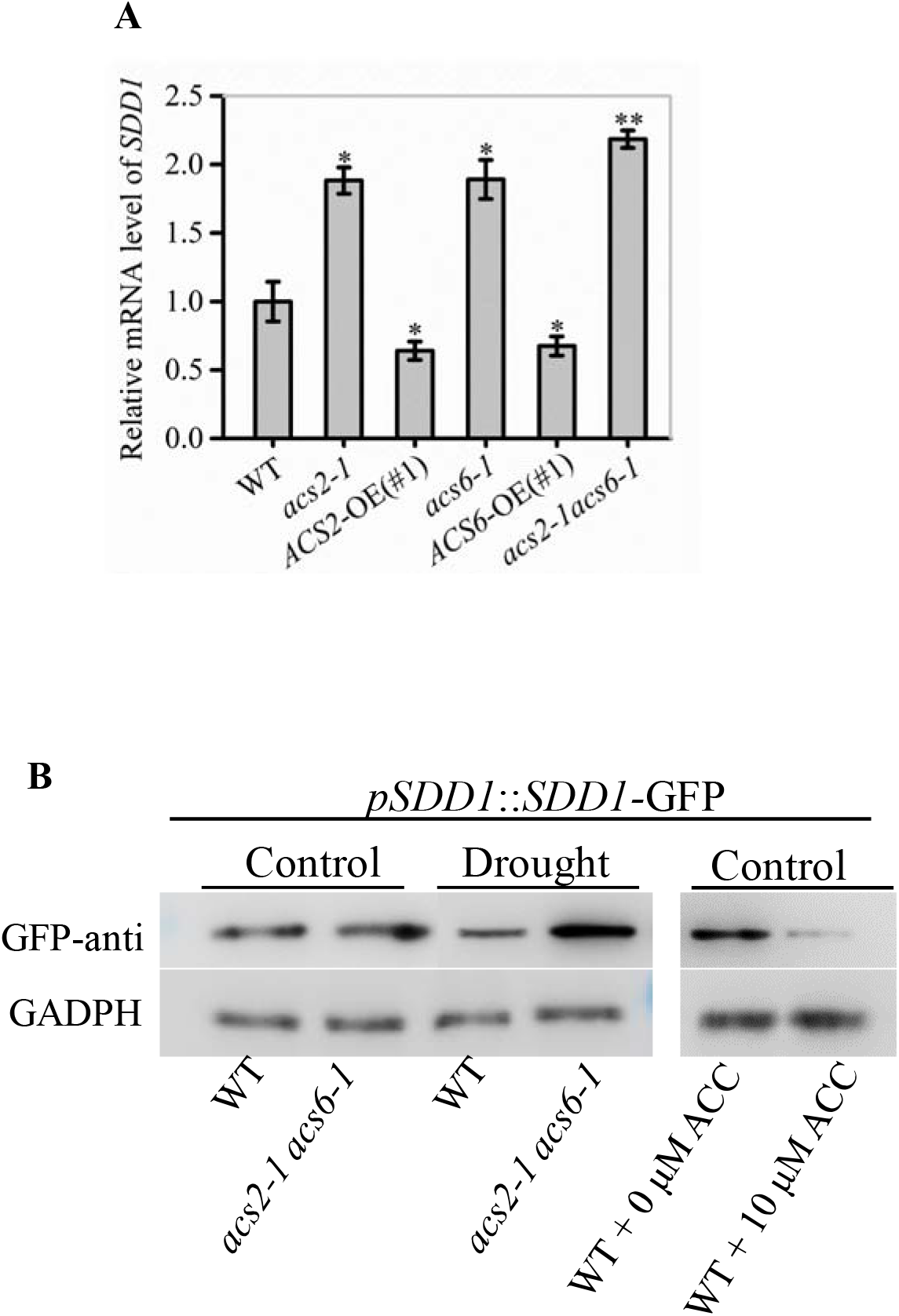
Inhibitory effects of ACS2/6 activation and ACC treatment on SDD1 expression and protein accumulation. (A) qRT-PCR-based relative levels of *SDD1* mRNA transcripts in immature leaves of WT, *acs2-1*, *acs6-1*, *acs2-1acs6-1*, *ACS2*-OE(#1), and *ACS6*-OE(#1) seedlings. The experiment was repeated three times with consistent results. Values are means ± SD (Student’s *t*-test; *, *P* < 0.05; **, *P* < 0.01). (B) Evaluation of SDD1 protein levels by western blotting. The fusion protein GFP-SDD1 was collected from immature leaves of *pSDD1*::*SDD1*-*GFP*-expressing WT or *acs2-1acs6-1* lines with or without drought and ACC (0 or 10 μM) treatment. Levels of GFP-SDD1 fusion protein were determined using GFP antibody. GAPDH protein was used as a loading control. The experiment was repeated three times with consistent results.

### SPCH activity positively regulated GTL1 expression

The trihelix transcription factor GTL1 is known to be the direct controller of SDD1 activity (Yoo *et al*., 2010; Weng *et al*., 2012; Virdi *et al*., 2015), and ChIP-sequencing data suggest that GTL1 is a target of SPCH (Lau *et al*., 2014). However, experimental evidence that SPCH affects GTL1 activity was lacking.

We investigated the effects of SPCH activity on *GTL1* expression. The *GTL1* mRNA levels in *spch-1*, *spch-3*, and *SPCH*-OE(#3) were, respectively, 0.43-, 0.36-, or 4.47-fold higher than that in WT (Fig. 7A). These data suggest that SPCH promoted *GTL1* expression. We verified this promoting effect by carrying out a transient transcription activity assay in tobacco leaves. Imaging analyses indicated that the presence of SPCH protein increased LUC fluorescence intensity linked to the specific promoter fragment of *GTL1* compared with that of the blank control, LUC alone (Fig. 7B). Interestingly, SPCH-stimulated LUC activity was 4.1-times higher than that of LUC alone (Fig. 7C), indicating that SPCH activated *GTL1* expression by binding to its promoter.

**Fig. 7.**
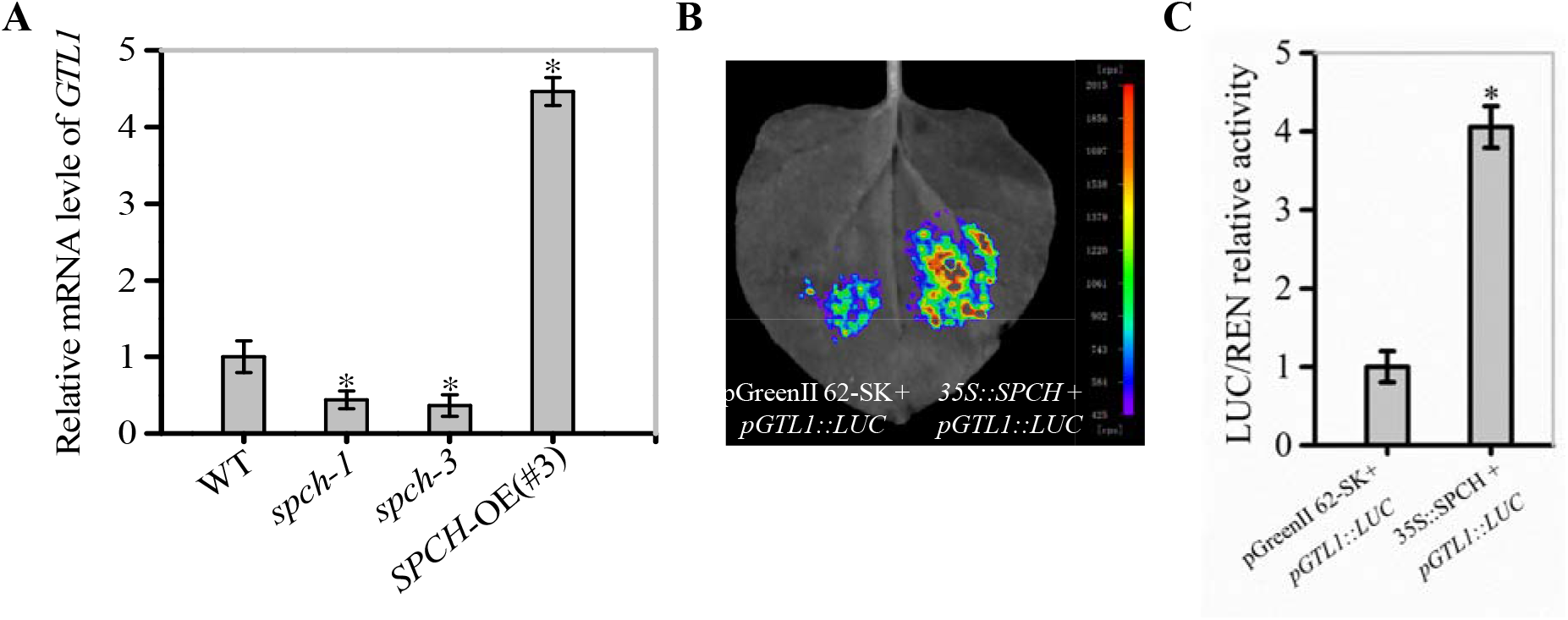
SPCH promotion of GTL1 expression. (A) qRT-PCR-based relative levels of *GTL1* mRNA transcripts in immature leaves of WT, *spch-1*, *spch-3*, and *SPCH*-OE(#3) plants. *ACTIN8* was used as an internal control. The experiment was repeated three times with consistent results. Values are means ± SD (Student’s *t*-test; *, *P* < 0.05). (B) and (C) Binding of SPCH protein to the *GTL1* promoter in tobacco leaves in a transient transcription dual-luciferase assay. The size and intensity of LUC fluorescence signals recorded by IndiGO software are proportional to binding ability (B). Relative binding ability was evaluated quantitatively by calculating the ratio of the fluorescence intensity of firefly luciferase (LUC) to that of an internal control, Renilla luciferase (REN) (C). Values are means ± SD (*n* = 3). Asterisks indicate significant differences (*, *P* < 0.05) compared with leaf regions injected with *Agrobacterium* harboring an empty vector.

### ACC buffered Ca^2+^ activity in stomatal lineage cells

Studies have shown that Ca^2+^-loaded CaM can relieve the inhibition of *SDD1* expression by GTL1, while the susceptibility of GTL1 to Ca^2+^ levels mainly occurs in stomatal lineage (Weng *et al*., 2012; Virdi *et al*., 2015; Yoo *et al*., 2019). Thus, we monitored how ACC mediates Ca^2+^ levels in stomatal lineage cells on the leaf epidermis. Using the Ca^2+^ fluorescence probe Fluo-4/AM, we preliminarily evaluated the Ca^2+^ levels in stomatal lineage cells on the immature leaf epidermis. Under normal conditions, the Ca^2+^ levels in stomatal lineage cells were slightly higher in *acs2-1acs6-1* than in WT. However, after halting watering for 6 days, the Ca^2+^ levels in stomatal lineage cells were higher in *acs2-1acs6-1* than in WT (Supplementary Fig. S5). This hints that the decreased ACC accumulation increased Ca^2+^ levels in stomatal lineage cells.

The Ca^2+^-sensitive yellow cameleon protein YC3.6 has been developed as a Ca^2+^ biosensor (Krebs *et al*., 2012; Behera *et al*., 2017). Therefore, we created NES-YC3.6 expressing *acs2-1acs6-1* lines to obtain *in vivo* data on Ca^2+^ accumulation in the stomatal lineage cells on the leaf epidermis. Both fluorescence-symbolized and cpVenus/CFP ratio-labelled Ca^2+^ levels were analyzed. Under normal watering conditions, the fluorescence intensity of YC3.6 protein was slightly higher in *acs2-1acs6-1* than in WT (Fig. 8A). Moreover, the cpVenus/CFP ratio was slightly higher in *acs2-1acs6-1* than in WT (Fig. 8B). After halting watering for 6 days, the fluorescence intensity of YC3.6 protein was significantly higher in *acs2-1acs6-1* than in WT (Fig. 8A). Specifically, the Ca^2+^ levels in the stomatal lineage cells was 3.38-times higher in *acs2-1acs6-1* than in WT (Fig. 8B). This result shows that ACC accumulation in leaves significantly reduced Ca^2+^ levels or activity in stomatal lineage cells.

**Fig. 8.**
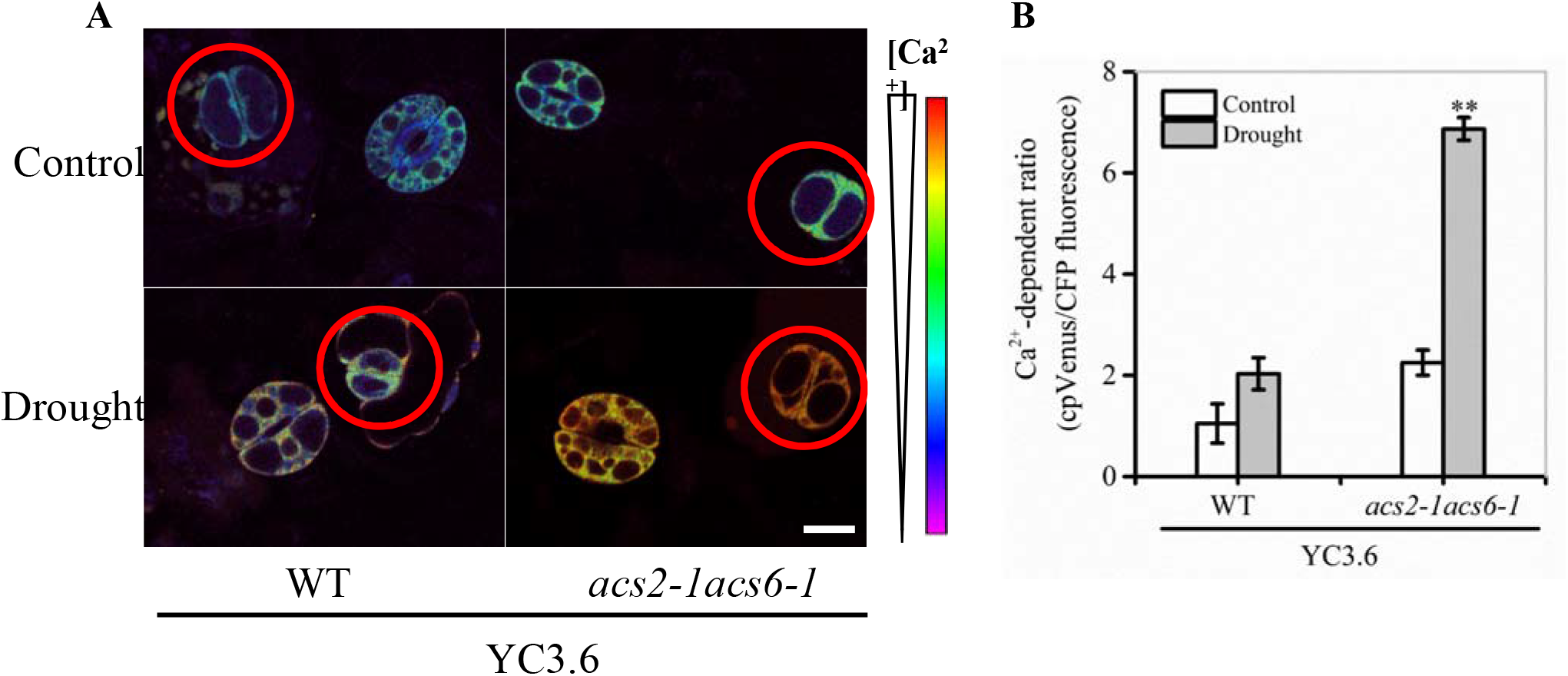
ACS2/6-generated ACC buffering of Ca^2+^ activity in stomatal lineage cells. (A) The relative fluorescence intensity of YC3.6 expression in stomatal lineage cells of immature leaves of NES-YC3.6-expressing WT or *acs2-1acs6-1* plants or by drought treatment. Representative images from at least 25 leaves in each line are shown. Pseudocolors in images correspond to relative Ca^2+^ levels according to the color scale on the right. Scale bar, 10 μm. (B) Evaluation of Ca^2+^ levels in stomatal lineage cells (labeled with red circle) of immature by calculating the ratio of the FRET acceptor cpVenus (at 525-555 nm) to the FRET donor CFP (at 465-499 nm). The experiment was repeated three times with consistent results. Values are means ± SD (Student’s *t*-test; **, *P* < 0.01).

## Discussion

These findings reveal the specific role of ACS2/6 activity in the establishment of stomatal density and pattern under drought. A schematic overview of the inter-relationships among these processes is provided in Fig. 9.

**Fig. 9.**
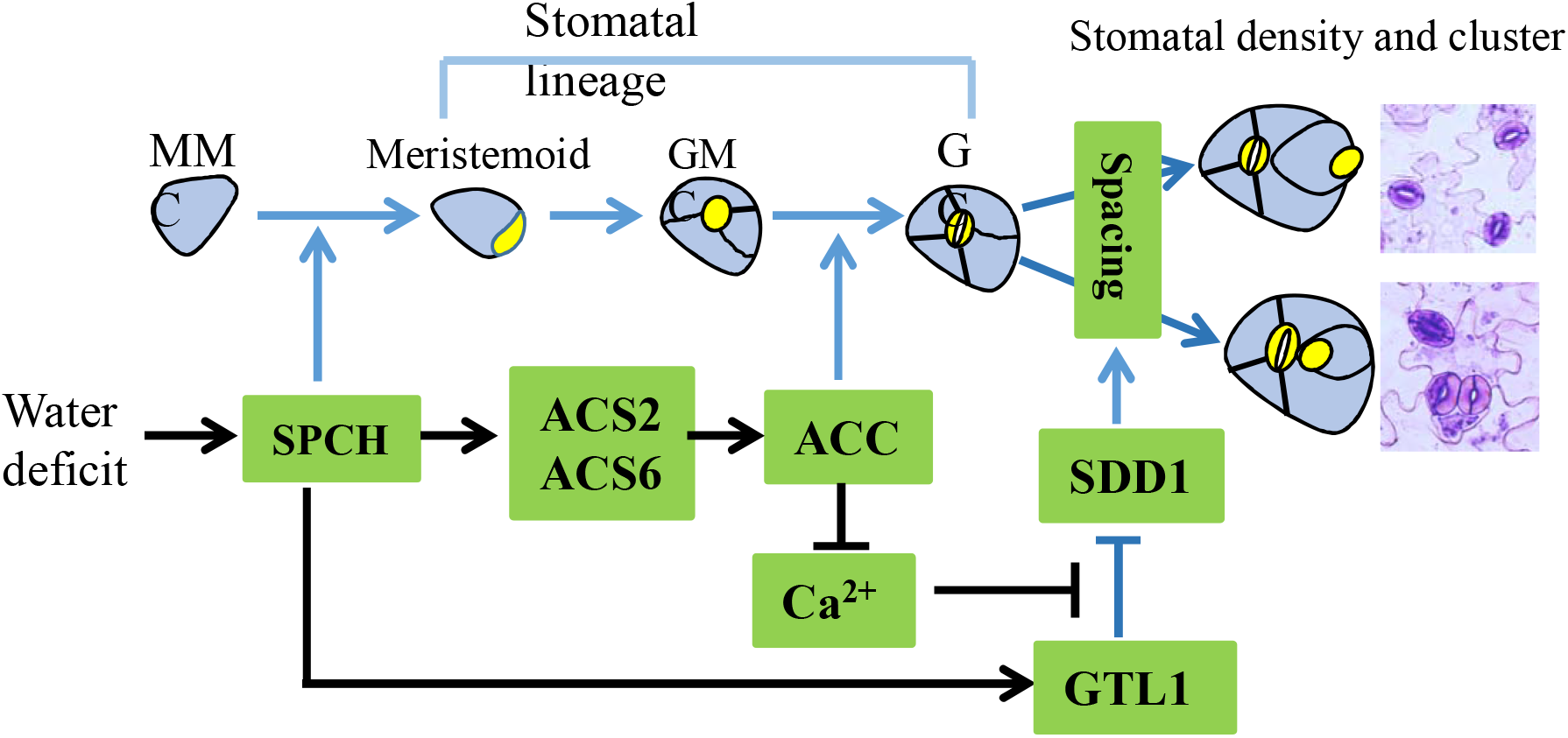
Diagram illustrating the role of ACS2/6 expression and ACC accumulation in the integration of SPCH-initiated stomatal development with SDD1-dependent stomatal spacing. In the top row depicting stomatal development, non-stomatal lineage cells, such as leaf epidermal cells, protodermal cells, and meristemoid mother cells (MMCs), are shown in blue, while stomatal lineage cells, including meristemoid cells, guard mother cells (GMCs), and guard cell (GCs), are indicated in yellow. Under water deficit conditions, SPCH increases ACS2, ACS6, and GTL1 expressions. Next, ACS2/6-generated ACC accumulation may be involved in two processes: 1) promoting the transformation of GMCs into GCs, and 2) inducing a shortage of Ca^2+^ in stomatal lineage cells and increasing SDD1 activity. As a consequence of these two processes, ACS2/6-generated ACC accumulation increases stomatal density and the rate of clustering, which ultimately leads to plant wilting and death under conditions of escalating drought. Relationships among these events are indicated by arrows: blue for conclusions drawn from the literature, and black for findings of the present study.

The activation of ACS2/6 lays the foundation for ACC-induced stomatal generation and pattern under drought. According to our data, stomatal density and clustering on the leaf epidermis were reduced in loss-of-function mutants *acs2-1*, *acs6-1*, and *acs2-1acs6-1*, but were increased in *ACS2-* and *ACS6-*overexpression lines (Fig. 3); to put it in another way, drought-activated ACS2/6 increased stomatal density and cluster, and these facilitated stomata-based water evaporation, in turn, seedlings withered and some even died with drought escalating. This finding provides a genetic explanation for the decrease in stomatal density and clustering caused by inhibitors of ACS activity, such as AVG or PAC, in *Arabidopsis* (Serna and Fenoll, 1996; Saibo *et al*., 2003; Yin *et al*., 2019), and also provide theoretical explanations why ACS2- or ACS6-deficient rice (Zhang *et al*., 2013) or maize (Young *et al*., 2004) are less sensitive to water deficit than are WT controls. Considering the specificity of ACS activity (Tsuchisaka and Theologis, 2004; Tsuchisaka *et al*., 2009; Han *et al*., 2019; Lv *et al*., 2019), we presume that ACS2/6 activation is specific to stomatal development and spacing on the leaf epidermis when *Arabidopsis* seedlings are under drought. Because drought can induce ACS2/6 activation and ACC accumulation (Catalá *et al*., 2014; Dubois *et al*., 2018), the observed increase in stomatal density and cluster under drought conditions (Fig. 2, 3) is easily understandable. Simply, ACS2/6-dependent ACC accumulation increases the susceptibility of seedlings to drought. Furthermore, activated ACS2 and ACS6 may function in parallel, as growth phenotypes (Supplementary Fig. S1), stomatal densities (Fig. 3B), and clustering (Fig. 3C) were similar among *acs2-1*, *acs6-1*, and *acs2-1acs6-1* mutants, and the expressions of *ACS6* and *ACS2* were relatively unaffected in the two single mutants *acs2-1* and *acs6-1*, respectively (Fig. 1B). Importantly, our results show that SPCH separately regulates the expression of *ACS2* and *ACS6* by binding to their promoters (Fig. 4). We speculate that ACS2 and ACS6 jointly ensure plants to fully respond to frequent drought stimuli.

ACS2/6-dependent ACC production is an important node of SPCH-based regulation of stomatal development and pattern. In line with a previous prediction (Lau *et al*., 2014), our results provide evidence that SPCH acts as a transcription factor to control the expression of *ACS2* and *ACS6* (Fig. 4). The ability of SPCH to promote *ACS2* and *ACS6* expression was evidenced by the fact, for example, that *spch-1* and *spch-3* mutants showed reduced expressions of these genes (Fig. 4) and ACC content, whereas SPCH overexpression led to increased ACC levels (Fig. 5). This finding explains why plant tolerance to osmotic stress requires reduced SPCH activity (Han and Torii, 2016; Tripathi *et al*., 2016; Zoulias *et al*., 2018).

ACS2/6-generated ACC accumulation acts as a bridge between SPCH-initiated stomatal individual development and SDD1-directed stomatal spacing between stomata. Evidence for this conclusion is as follows: First, ACC mimics SPCH to reduce SDD1 activity, thereby increasing stomatal density and cluster. The mutants *acs2-1*, *acs6-1*, and *acs2-1acs6-1* seedlings (Fig. 2) mimicked transgenic *SDD1-*overexpressing plants by showing reduced stomatal density and cluster on leaves. Consistent with this observation, both the *sdd1-1* mutant (Von *et al*., 2002) and ACS2*-* and ACS6*-*overexpressing plants (Fig. 6) exhibited increased stomatal density and cluster on the leaves. Second, ACC-associated Ca^2+^ insufficiency reduced SDD1 activity, or, alternatively, Ca^2+^ activity, in stomatal lineage cells, so that ACC levels were linked to SDD1 expression. Our findings indicated that Ca^2+^ levels in stomatal lineage cells on the leaf epidermis were higher in *acs2-1acs6-1* plants than in WT (Fig. 8; Supplementary Fig. S5). This suggests that ACC accumulation inhibits SDD1 activity by controlling Ca^2+^ activity in stomatal lineage cells. This result is reasonable because a Ca^2+^ shortage can stabilize the binding of GTL1 to the *SDD1* promoter to prevent its expression in stomatal lineage cells (Yoo *et al*., 2019). These findings explicate the mechanisms in the recent discovery that Ca^2+^ activity intensifies stomata-based water evaporation from leaves of *Arabidopsis* seedlings under drought conditions (Teardo *et al*., 2019).

In brief, these findings first indicated that the ACS2/6 activity may specifically integrated SPCH-initiated stomatal individual development with SDD1-directed space setting between stomata under drought. This integration increased stomatal density and cluster on the leaf epidermis under moderate drought, which laid foundation for seedling wilting and death with drought escalating. The promotion of moderate drought to stomatal density and cluster provided a hint that the evolutionary memory of plants from aquatic to terrestrial may be evoked (Croxdale, 2000); while this evolutionary memory appears to override routine terrestrial regulation determining stomatal development (Croxdale, 2000; van Veen and Sasidharan, 2021).

## Acknowledgements

This work was supported by the National Natural Science Foundation of China (31970735, 31271510) to Jing Jiang. We thank Professor Xiao-lan Chen (School of Life Sciences, Yunnan University, China), Professor Sui-wen Hou (MOE Key Laboratory of Cell Activities and Stress Adaptations, Lanzhou, China), and Professor Jörg Kudla (Institute of Biology and Biotechnology of Plants, University of Munster, Germany) provided seeds of *spch-1*, *spch-3*, the transgenic *pSPCH*::*SPCH*-*GFP* line and the Ca^2+^ sensor NES-YC3.6-expressing line, respectively.

## Authors’ contributions

JJ and CPS formulated the experimental strategy. MZJ, JJ, LYL and CG performed experiments. JJ and MZJ wrote the paper.

## Data availability

All data relevant to this study are presented in figures and supplementary data.

## Supplementary Data

Table. Primer list.

Fig. S1. ACS2- or ACS6-based growth phenotype of *Arabidopsis* seedling or with drought treatment.

Fig. S2. ACC-increased stomatal density and percentage of clustering stomata on leaf epidermis.

Fig. S3. The control experiment of SPCH-activated ACS2 or ACS6 expression, respectively, in the transient transcription dual-luciferase assay system.

Fig. S4. RT-qPCR analysis on the inhibitory effect of ACC treatment on the expression activity of *SDD1* gene in immature leaves.

Fig. S5. Ca^2+^-stimulated Fluo-4/AM fluorescence intensity in stomatal lineage cells on leaf epidermis.

## Notes

### Competing Interest Statement

The authors have declared no competing interest.

## References

Acharya BR, Assmann SM. 2009. Hormone interactions in stomatal function. Plant Molecular Biology 69, 451–462.

Bao Y, Wang C, Jiang C, Pan J, Zhang G, Liu H, Zhang H. 2014. The tumor necrosis factor receptor-associated factor (TRAF)-like family protein SEVEN IN ABSENTIA 2 (SINA2) promotes drought tolerance in an ABA-dependent manner in *Arabidopsis*. New Phytologist 202, 174–187.

Behera S, Long Y, Schmitz-Thom I, et al. 2017. Two spatially and temporally distinct Ca^2+^ signals convey *Arabidopsis thaliana* responses to K^+^ deficiency. New Phytologist 213, 739–750.

Berger D, Altmann T. 2000. A subtilisin-like serine protease involved in the regulation of stomatal density and distribution in *Arabidopsis thaliana*. Genes Development 14, 1119–1131.

Bleecker AB, Kende H. 2000. Ethylene: a gaseous signal molecule in plants. Annual Review of Cell and Developmental Biology 16, 1–18.

Casson S, Gray JE. 2008. Influence of environmental factors on stomatal development. New Phytologist 178, 9–23.

Catalá R, López-Cobollo R, Mar Castellano M, Angosto T, Alonso JM, Ecker JR, Salinas J. 2014. The *Arabidopsis* 14-3-3 protein RARE COLD INDUCIBLE 1A links low-temperature response and ethylene biosynthesis to regulate freezing tolerance and cold acclimation. Plant Cell 26, 3326–3342.

Croxdale JL. 2000. Stomatal patterning in angiosperms. American Journal of Botany 87, 1069–1080.

de Marcos A, Houbaert A, Triviño M, Delgado D, Martín-Trillo M, Russinova E, Fenoll C, Mena M. 2017. A mutation in the bHLH domain of the SPCH transcription factor uncovers a BR-dependent mechanism for stomatal development. Plant Physiology 174, 823–842.

Dubois M, Claeys H, Van den Broeck L, Inzé D. 2017. Time of day determines *Arabidopsis* transcriptome and growth dynamics under mild drought. Plant Cell Environment 40, 180–189.

Dubois M, Van den Broeck L, Inzé D. 2018. The pivotal role of ethylene in plant growth. Trends Plant Science 23, 311–323.

Hamanishi ET, Thomas BR, Campbell MM. 2012. Drought induces alterations in the stomatal development program in *Populus*. Journal of Experimental Botany 63, 4959–4971.

Han S, Jia MZ, Yang JF, Jiang J. 2019. The integration of ACS2-generated ACC with GH3-mediated IAA homeostasis in NaCl-stressed primary root elongation of *Arabidopsis* seedling. Plant Growth Regulation 88, 151–158.

Han SK, Torii KU. 2016. Lineage-specific stem cells, signals and asymmetries during stomatal development. Development 143, 1259–1270.

Hepworth C, Doheny-Adams T, Hunt L, Cameron DD, Gray JE. 2015. Manipulating stomatal density enhances drought tolerance without deleterious effect on nutrient uptake. New Phytologist 208, 336–341.

Krebs M, Held K, Binder A, Hashimoto K, Den Herder G, Parniske M, Kudla J, Schumacher K. 2012. FRET-based genetically encoded sensors allow high-resolution live cell imaging of Ca^2+^ dynamics. The Plant Journal 69, 181–192.

Lau OS, Davies KA, Chang J, Adrian J, Rowe MH, Ballenger CE, Bergmann DC. 2014. Direct roles of SPEECHLESS in the specification of stomatal self-renewing cells. Science 345, 1605–1609.

Livak KJ, Schmittgen TD. 2001. Analysis of relative gene expression data using real-time quantitative PCR and the 2^−ΔΔC(T)^ method. Methods 25, 402–408.

Lv SF, Jia MZ, Zhang SS, Han S, Jiang J. 2019. The dependence of leaf senescence on the balance between 1-aminocyclopropane-1-carboxylate acid synthase 1 (ACS1)-catalyzed ACC generation and nitric oxide associated 1 (NOS1)-dependent NO accumulation in *Arabidopsis*. Plant Biology 21, 595–603.

MacAlister CA, Ohashi-Ito K, Bergmann DC. 2007. Transcription factor control of asymmetric cell divisions that establish the stomatal lineage. Nature 445, 537–540.

Pillitteri LJ, Sloan DB, Bogenschutz NL, Torii KU. 2007. Termination of asymmetric cell division and differentiation of stomata. Nature 445, 501–505.

Qi SL, Lin QF, Feng XJ, Han HL, Liu J, Zhang L, Wu S, Le J, Blumwald E, Hua XJ. 2019. IDD16 negatively regulates stomatal initiation via trans-repression of SPCH in *Arabidopsis*. Plant Biotechnol Journal 17, 1446–1457.

Saibo NJ, Vriezen WH, Beemster GT, Van Der Straeten D. 2003. Growth and stomata development of *Arabidopsis* hypocotyls are controlled by gibberellins and modulated by ethylene and auxins. Plant Journal 33, 989–1000.

Serna L, Fenoll C. 1996. Ethylene induces stomata differentiation in *Arabidopsis*. International Journal of Developmental Biology 1, 123S–124S.

Serna L, Fenoll C. 1997. Tracing the ontogeny of stomatal clusters in *Arabidopsis* with molecular markers. Plant Journal 12, 747–755.

Serna L. 2009. Cell fate transitions during stomatal development. Bioessays 31, 865–873.

Teardo E, Carraretto L, Moscatiello R, et al. 2019. A chloroplast-localized mitochondrial calcium uniporter transduces osmotic stress in *Arabidopsis*. Nature Plants 5, 581–588.

Tripathi P, Rabara RC, Reese RN, et al. 2016. A toolbox of genes, proteins, metabolites and promoters for improving drought tolerance in soybean includes the metabolite coumestrol and stomatal development genes. BMC Genomics 17, 102.

Tsuchisaka A, Theologis A. 2004. Unique and overlapping expression patterns among the *Arabidopsis* 1-amino-cyclopropane-1-carboxylate synthase gene family members. Plant Physiology 136, 2982–3000.

Tsuchisaka A, Yu G, Jin H, Alonso JM, Ecker JR, Zhang X, Gao S, Theologis A. 2009. A combinatorial interplay among the 1-aminocyclopropane-1-carboxylate isoforms regulates ethylene biosynthesis in *Arabidopsis thaliana*. Genetics 183, 979–1003.

van Veen H, Sasidharan R. 2021. Shape shifting by amphibious plants in dynamic hydrological niches. New Phytologist 229, 79–84.

Virdi AS, Singh S, Singh P. 2015. Abiotic stress responses in plants: roles of calmodulin-regulated proteins. Frontiers in Plant Science 6, 809.

Von Groll U, Berger D, Altmann T. 2002. The subtilisin-like serine protease SDD1 mediates cell-to-cell signaling during *Arabidopsis* stomatal development. Plant Cell 14, 1527–1539.

Weng H, Yoo CY, Gosney MJ, Hasegawa PM, Mickelbart MV. 2012. Poplar GTL1 is a Ca^2+^/calmodulin-binding transcription factor that functions in plant water use efficiency and drought tolerance. PLoS One 7, e32925.

Xie Z, Nolan T, Jiang H, Tang B, Zhang M, Li Z, Yin Y. 2019. The AP2/ERF transcription factor tiny modulates brassinosteroid-regulated plant growth and drought responses in *Arabidopsis*. Plant Cell 31, 1788–1806.

Yin J, Zhang X, Zhang G, Wen Y, Liang G, Chen X. 2019. Aminocyclopropane-1-carboxylic acid is a key regulator of guard mother cell terminal division in *Arabidopsis thaliana*. Journal of Experimental Botany 70, 897–908.

Yoo CY, Mano N, Finkler A, Weng H, Day IS, Reddy ASN, Poovaiah BW, Fromm H, Hasegawa PM, Mickelbart MV. 2019. A Ca^2+^/CaM-regulated transcriptional switch modulates stomatal development in response to water deficit. Scientific Reports 9, 12282.

Yoo CY, Pence HE, Jin JB, Miura K, Gosney MJ, Hasegawa PM, Mickelbart MV. 2010. The *Arabidopsis* GTL1 transcription factor regulates water use efficiency and drought tolerance by modulating stomatal density via transrepression of SDD1. Plant Cell 22, 4128–4141.

Young TE, Meeley RB, Gallie DR. 2004. ACC synthase expression regulates leaf performance and drought tolerance in maize. Plant Journal 40, 813–825.

Zhang H, Zhang J, Quan R, Pan X, Wan L, Huang R. 2013. EAR motif mutation of rice OsERF3 alters the regulation of ethylene biosynthesis and drought tolerance. Planta 237, 1443–1451.

Zoulias N, Harrison EL, Casson SA, Gray JE. 2018. Molecular control of stomatal development. Biochemical Journal 475, 441–454.

